# Phylo-MIP: Phylogeny-based Molecular Identification Pipeline for DNA metabarcoding, and assessment of insect communities in subalpine river ecosystems

**DOI:** 10.1101/2025.11.10.687572

**Authors:** Takumi Yshida, Shonosuke Shigeta, Yuta Hasebe, Masaki Takenaka

## Abstract

Environmental DNA metabarcoding is a powerful tool for biodiversity assessment; however, species identification often relies on sequence similarity searches (BLAST), which are limited by incomplete reference databases and result in frequent misidentifications or unclassified taxa. To overcome these challenges, we designed a novel analytical pipeline, Phylo-MIP, which integrates phylogenetic analysis and species delimitation. Phylo-MIP combines sequence alignment, haplotype thinning, phylogenetic tree construction, and multiple species delimitation methods (bPTP, bPTP-ML, mPTP) to objectively identify Molecular Operational Taxonomic Units (MOTUs) without relying on reference databases. As a validation, we applied Phylo-MIP to previously published eDNA data from river in Kanagawa Prefecture, detecting approximately 28% more species than only BLAST-based methods. We also conducted a eDNA survey in the subalpine zone of Kamikochi, Nagano Prefecture, Japan, where Phylo-MIP revealed distinct community structures associated with specific aquatic environments. Phylo-MIP enables non-invasive and efficient biodiversity monitoring, even in protected areas, and offers a reproducible framework for ecological research. This approach has the potential to become a new standard method in river ecology, taxonomy, conservation biology, and ecosystem management.

## 1 Introduction

Understanding the biota provides the most basic information in ecology and evolutionary biology (Kelly et al., 2016; Deiner et al. 2017). Moreover, for environmental conservation and management, species richness is essential for developing strategies. However, obtaining biological information and conducting biological monitoring involve various issues and significant costs. Additionally, understanding biota, including rare species, often requires partial environmental destruction and faces taxonomic problems (Kelly et al., 2016). Metabarcoding analysis, especially environmental DNA (eDNA) monitoring, is a fast and efficient method to survey species richness in various fields (Miya et al. 2015; Creer et al., 2016; Deiner et al., 2017). Several advantages of eDNA metabarcoding include its higher cost and time effectiveness, its noninvasive nature, and its high specificity and sensitivity (Burian et al. 2021). Additionally, by analyzing materials such as animal feces (Milner et al. 2020; Nagahara et al. 2025) or the stomach contents of fish (Jo et al. 2016; Siegenthaler et al. 2019), it is possible to determine what target species have consumed. Particularly with the use of next-generation sequencing (NGS), eDNA monitoring becomes a powerful tool that is expected to advance further in the future. Metabarcoding involves amplifying specific DNA barcoding regions from a sample and comprehensively obtaining DNA reads (nucleotide sequences) using NGS.

However, the process of referencing obtained sequences against a database to identify species is not something that can be easily performed by everyone. It requires a certain level of bioinformatics skills and specialized knowledge of biological species identification. Molecular species identification typically involves using DNA barcoding regions specific to the target group Traditional methods for calculating ecological indices from metabarcoding data typically involve clustering DNA reads into operational taxonomic units (OTUs), followed by taxonomic assignment using reference databases (e.g., Kermarrec et al., 2014; Zimmermann et al., 2015). Species identification is often performed by comparing sequence similarity between reads and reference sequences using online tools such as NCBI (National Center for Biotechnology Information) BLAST (Altschul et al., 1997), where species determination is based on sequence similarity, often using thresholds like 98% or 97% (Uchida *et al*. 2020; Sander *et al*. 2024; Takenaka et al. 2024). Despite its widespread use, this approach faces several limitations. One major issue is that many OTUs remain unclassified at the species level due to gaps in reference libraries (Keck et al., 2018; Takenaka et al. 2025). Furthermore, the resulting taxonomic assignments generally lack integration with phylogenetic frameworks (Czech et al., 2022). When the sequence similarity exceeds a certain threshold (e.g., 2, 3%), the sequences are treated as distinct Molecular Operational Taxonomic Units (MOTUs). However, there is no particular reason to assume that individuals from different species share the same average level of similarity, nor is there a clear justification for setting the threshold specifically at 3% (De Santiago et al. 2022; Diniz-Filho et al. 2024). Several methods have been proposed to incorporate phylogenetic interpretation by placing genetic sequences onto a reference phylogenetic tree (e.g., EPA-NG [Barbera et al., 2019], PPLACER [Matsen et al., 2010], and phyloBARCODER [Inoue et al., 2024]). However, interpreting the results of phylogenetic analyses objectively requires advanced expertise.

We constructed methods of a molecular species identification that incorporates phylogenetic interpretation and species delimitation analyses. Phylogenetic analysis allows genetic similarities to be recognized as clusters (so-called genetic lineages or MOTUs, de novo species). By using phylogenetic analysis to identify these clusters as units of species, more accurate results can be obtained compared to relying solely on sequence similarity thresholds, even when dealing with incomplete DNA databases or genetically similar species. Similar approaches have been proposed and examined in previous studies. However, even if phylogenetic analysis is conducted, accurately interpreting phylogenetic results can sometimes be challenging. This difficulty can persist even for experts, who may find it hard to provide objective interpretations in certain cases. Therefore. our purpose is developing a method that enables consistent interpretation by anyone. We implemented species delimitation analyses, commonly used in phylogenetic studies, to establish an objective framework for species separation based on defined criteria. By applying these analyses, we were able to assign MOTU or de novo species-level information directly to sequence data, thereby reducing reliance on reference databases such as BLAST. This approach also helps avoid underestimation of species richness, which is often a limitation in database-dependent identification methods (Fig. 1).

**Figure 1.**
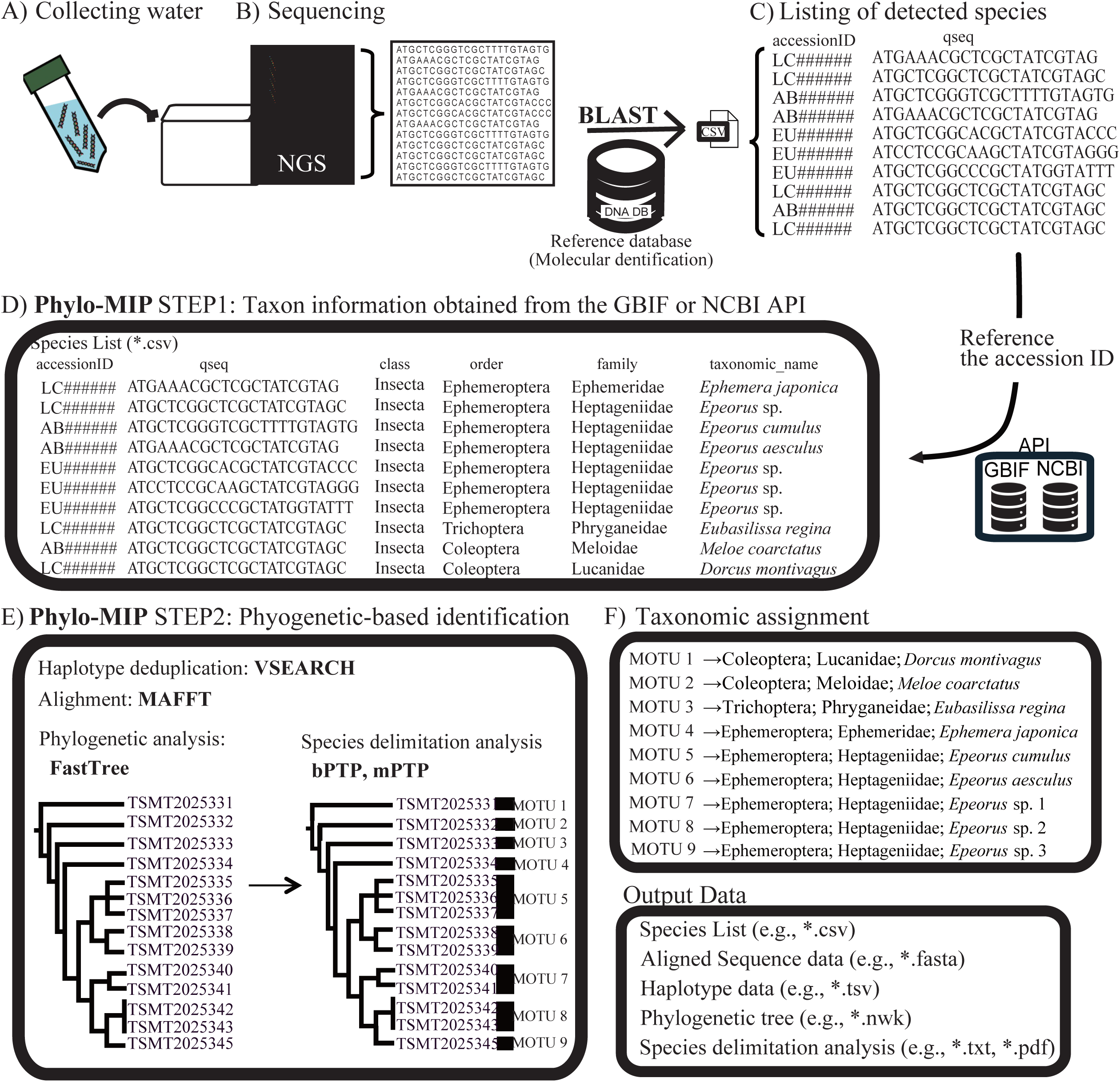
Overview of the Phylo-MIP workflow for metabarcoding DNA analysiss: A) Collecting water samples; (B) Sequencing: eDNA metabarcoding is employed using next-generation sequencing (NGS); (C) Listing detected species: Raw sequences are processed, and taxonomic identification is initially performed using BLAST search against reference databases; (D) Phylo-MIP Step 1 – Taxonomic information retrieval: BLAST results are used to obtain taxonomic information via GBIF and NCBI APIs, generating a species list with hierarchical classification (class, order, family, species); (E) Phylo-MIP Step 2 – Phylogenetic-based identification: Sequences are aligned (MAFFT), haplotypes are deduplicated (VSEARCH), and phylogenetic trees are constructed (FastTree). Species delimitation analyses (bPTP, bPTP-ML, mPTP) are applied to define Molecular Operational Taxonomic Units (MOTUs); (F) Taxonomic assignment and output: final data.

In this study, we first conducted a BLAST search of the obtained sequences, following the conventional method of manually identifying species based on a 98% similarity threshold. Subsequently, we performed phylogenetic analysis and species delimitation analysis to propose an objective and standardized method for genetic species identification that can be consistently applied by anyone. To establish our method, we focused on aquatic insects in Kamikochi. Kamikochi is located in the subalpine zone of the Japanese Islands and is fed by mountain ranges exceeding 3,000 meters in elevation. As a result, it features various aquatic environments, including springs and wetlands. Kamikochi has long been designated as a national park and protected, yet comprehensive surveys of insects have not been conducted. Regarding aquatic insects, the only study has been by Nozaki et al. (2014), which focused on caddisflies. Therefore, we collected water samples from various environments in Kamikochi, including the main river, tributaries, wetlands, and springs, to investigate whether aquatic insects inhabit these areas and to explore trends in species distribution across different environments.

## 2 Materials and Methods

In this study, we evaluated the accuracy and detection efficiency of species identification using phylogenetic approaches in eDNA metabarcoding analyses, based on the eDNA data obtained by Takenaka et al. (2024) which conducted eDNA analyses at six sites in the two river systems in Japanese Islands (the Sagami-gawa River and Sakawa-gawa River systems; 3 replicates per site). Then, we conducted an eDNA based survey of aquatic insect communities in Kamikochi which is a nationally designated cultural property and part of Chubu-Sangaku National Park using the method established in this study.

### 2.1 Pipeline Workflow

The data used for the analyses were obtained from a previous study (DDBJ: PRJDB15936; Takenaka et al., 2024). The purpose of using phylogenetic approaches for species identification was to identify and cluster OUT (Operational Taxonomic Unit)/ASV (Amplicon Sequence Variant) data derived from metabarcoding analyses. Specifically, the purpose was to assign molecular operational taxonomic unit (MOTU) information to the OTU/ASV data obtained from metabarcoding analyses and then conduct identification and clustering. We employed species delimitation analyses based on models such as the Poisson Tree Processes (PTP) model for identification and clustering. These approaches do not rely on taxonomic information, instead identifying MOTUs directly from OTU/ASV data (Floyd et al., 2002; Blaxter & Floyd, 2003; Goldstein & DeSalle, 2011). Unlike methods that rely on sequence similarity–based species identification, such as BLAST searches, these methods treat species as MOTUs based on phylogenetic analyses and species delimitation approaches. This allows for the provisional assignment of MOTU classes, even for taxa whose sequences do not match existing DNA database entries, which is often the case due to limited taxonomic research. Thus, a more accurate assessment of biodiversity is enabled, including α-diversity within sites and β-diversity among sites (Valentini et al., 2009). To facilitate analyses without requiring prior knowledge of phylogenetic methods, we design a pipeline named “Phylogeny-based Molecular Identification Pipeline (Phylo-MIP)” (https://github.com/shigebio/Phylo-MIP: Fig. 1). This pipeline enables taxonomic assignment of OTU/ASV data using a complementary approach with representative MOTU sequences. We adopted the PTP model for MOTU identification for several reasons. As an analysis based on phylogenetic tree information, it allows MOTU identification while capturing the evolutionary history of the target group simultaneously. Additionally, its extensions, multi-rate PTP (mPTP) (Kapli et al., 2017) and Bayesian PTP (bPTP) (Zhang et al., 2013), provide integrated visual information combining phylogenetic relationships and species delimitation. This facilitates interpretation and complementary insights based on user knowledge, such as morphological traits, genetic distances between closely related species, or phylogeographic information. The application of phylogeny-based species delimitation methods to large datasets is often limited by computational constraints (Puillandre et al., 2021). However, “Phylo-MIP” is suitable for analyses using small- to medium-sized datasets, such as those focusing on specific taxa at the order or family level within a given region. Additionally, while conventional phylogenetic analyses require expert knowledge to define species boundaries, PTP-based species delimitation provides a framework that is objective, reproducible, and accessible to any user.

Our pipeline (Phylo-MIP) consists of the following four processes (showed Fig. 1). Development and testing were performed using Python 3.7.15 (default, Mar 7 2025, 07:25:12). To ensure reproducibility and manage dependencies, the environment was containerized using the platform Docker version 28.0.4 (https://www.docker.com), which mitigates conflicts with other software and prevents variations caused by the host computer system (Grüning et al., 2018). The base image selected was Ubuntu 20.04 (https://ubuntu.com), which is capable of supporting all software components used in this pipeline.

### 2.2 Identification of taxonomic information

When a researcher performs a BLAST search against a local database constructed from sequence information obtained from NCBI, the Phylo-MIP can assign taxonomic information to OTU/ASV data based on accession IDs. The assignment of taxonomic group information is performed according to the following flow. (1) Phylo-MIP obtains the species name by executing an API request to the NCBI database. This step utilizes the ‘efetch’ function of Biopython’s Entrez module (Cock et al., 2009). The request key is the ‘sallac’ value, which corresponds to the NCBI accession ID included in the BLAST search results. (2) The pipeline executes a request to the GBIF Species API (https://techdocs.gbif.org/en/openapi/v1/species), using the species name obtained in the previous step as the key. Subsequently, the taxonomic category information is extracted from the response JSON data. In this process, we do not utilize the taxonomic information registered with NCBI. This is because the data structure of NCBI’s taxonomic information makes it difficult to mechanically and appropriately extract taxonomic data. However, if taxon information could not be obtained from GBIF (the Global Biodiversity Information Facility), an API request is sent to NCBI using the accession ID as the key to obtain taxon information that may be uncertain. The source of taxonomic group information is recorded in the ‘sourcè column of the *.csv file listing taxonomic group information for each OTU/ASV data. The obtained taxonomic groups are assigned as MOTUs under the following rules and used for subsequent phylogenetic analysis depending on the ‘pident’ value. Taxonomic category application rule: (a) Species name, pident >= 98.00; (b) Genus name, 95.00 <= pident < 98.00; (c) Family name, 90.00 <= pident < 95.00; (d) Order name, 85.00 <= pident < 90.00; (e) pident < 85.00: Assign the string “Low_Identity_Match”. Furthermore, for databases constructed from data obtained from NCBI, the ‘sallac’ value is assigned as the name for the OTU/ASV data. Data with low match rates at the time of BLAST search or inaccurate assigned taxonomic information may have been misidentified, so they were visually inspected and corrected as necessary.

Phylo-MIP workflow is as follows.

1. Align sequence

The aligned portion of the query sequence (qseq) output from BLAST is extracted to generate a *.fasta file. An alignment is then performed on the generated *.fasta file using MAFFT v7.453 (Katoh and Standley, 2013).

2. Identify haplotypes and thin out redundant haplotypes

Within the same region and location, species with differing degrees of genetic diversity coexist (Baggiano et al. 2010). Therefore, OTU/ASV data obtained from eDNA metabarcoding analysis are likely to include multiple species with differing genetic diversity, and the number of OTUs/ASVs detected per species is expected to be unevenly distributed across MOTUs. When conducting species delimitation analyses using tree files composed of such imbalanced datasets (subtrees with a large number of extremely short branches), the interpretation of MOTU identification requires caution (Zhang et al. 2013). In the Phylo-MIP, to reduce the impact of such imbalanced datasets on phylogenetic analysis, we employed a process of identifying haplotypes within OTU/ASV data and then thinning to ensure one sequence per haplotype. Haplotype identification and thinning of identical haplotypes were performed using VSEARCH v2.14.1 (Rognes et al., 2016) on sequence data aligned by MAFFT.

3. Phylogenetic analysis

Phylogenetic analyses are performed using FastTree version 2.1.11 (Price et al., 2009, 2010), which allows for the choice of maximum likelihood or neighbor-joining methods, is compatible with large data sets, and is fast to process.

4. Species delimitation alaysis

Species delimitation analysis based on the PTP model is performed on the tree file (*.nwk) output from FastTree. In species delimitation analysis, it is known that the MOTUs identified by different methods can differ even for the same dataset. Therefore, genetic species are usually determined by comparing the analysis results output by multiple methods (Tang et al. 2014; Blair & Bryson Jr. 2017; Luo et al. 2018; Song et al. 2018; Siriwut et al. 2021; Zarei et al. 2021). Our pipeline uses three methods to identify MOTUs: bPTP (Zhang et al. 2013), bPTP-ML (Zhang et al. 2013), and mPTP (Kapli et al. 2017).

Phylo-MIP outputs the following data:

- Sequence data (*.fasta) before and after alignment.
- Haplotype identification results and sequence data thinned to one sequence per haplotype (*.fasta)
- Phylogenetic analysis results (*.nwk)
- Results of each species delimitation analysis (image and text files).
- A CSV file that integrates the results of BLAST searches for each OTU/ASV, taxon information obtained from the GBIF or NCBI API, and the results of species delimitation analysis.

These allow quick and easy identification of OTU/ASV data obtained from metabarcoding analysis to taxa and addition of MOTU information. The document was developed to assist users in applying “Phylo-MIP” (https://github.com/shigebio/Phylo-MIP/blob/main /README.md).

### 2.3 Environmental DNA metabarcoding using Phylo-MIP in Kamikochi

At each site, 2 L of surface water was collected and filtered for eDNA analyses using Sterivex (0.45 μm pore size, Merck & Co., Inc.) in Kamikochi. Then, the filters were stored a at -20°C until eDNA analyses. Then, the filters were purified using a DNeasy Blood and Tissue kit (Qiagen). Finally, the column was eluted with 100 μL of AE buffer. Purified DNA were outsourced to Bioengineering Lab. Co., Ltd. They used the DNeasy PowerClean Pro Cleanup Kit (QIAGEN) to remove PCR inhibitors. DNA concentrations were measured using the Synergy LX microplate reader (Agilent Technologies) in combination with the QuantiFluor dsDNA System (Promega). Library preparation was performed using a 2-step tailed PCR method.

The first PCR for samples No. 1–14 was conducted using ExTaq HS (TaKaRa) with primers containing adapter sequences, MtInsects-16S_F (5′-GGA CGA GAA GAC CCT WTA GA-3′) and MtInsects-16S_R (5′-AT CCA ACA TCG AGG TCG CAA-3′), to assign the adapter region for the second PCR (Takenaka et al., 2023). For samples No. 23–26, modified primer sequences were used, namely AQdb16S_F (5′–TRA CYG TRC AAA GGT AGC–3′) and AQdb16S_R (5′– CCG GTY TRA ACT CAR ATC ATG T–3′). Each reaction was carried out in a 10 μL volume containing 1.0 μL of 10× Ex Buffer, 0.8 μL of dNTPs, 0.075 μL of ExTaq HS, 10 μM of each primer, and 2 μL of DNA template. Eight replicates were performed per sample, and the products from these eight reactions were pooled and used as the template for the second PCR. The thermal cycling conditions were as follows: initial denaturation at 94°C for 2 min; 35 cycles of 94°C for 30 s, 50°C for 30 s, and 72°C for 30 s; followed by a final extension at 72°C for 5 min. PCR products were purified using VAHTS DNA Clean Beads (Vazyme) at a 1.0× ratio to the reaction volume. The second PCR was performed using KOD FX Neo (TOYOBO). PCR products were again purified using VAHTS DNA Clean Beads (Vazyme) at a 1.0× ratio to the reaction volume. Sequencing was performed using either the NextSeq 1000 system with the NextSeq 1000/2000 P1 Reagents (600 Cycles) kit (Illumina) or the MiSeq system with the MiSeq Reagent Kit v3 (Illumina), under paired-end 2×300 bp conditions.

#### Bioinformatic analysis

Quality control of the FASTQ data was performed using fastp (Chen et al. 2018). The options specified were removal of sequences with a quality score below Q-Score30 (-q 30) and removal of sequences shorter than 50 bp (-l 50), with all other parameters kept at their default settings. After quality control, QIIME2 (Bolyen et al. 2019) was used with the cutadapt plugin to trim the PCR primers (--p-front-f GGACGAGAAGACCCTWTAGA / --p-front-r ATCCAACATCGAGGTCGCAA). The --p-discard-untrimmed option was enabled to remove reads in which primer sequences were not recognized. Subsequently, DADA2 (Callahan et al. 2016) was used for denoising and correction of sequencing errors. Because the 16S rRNA region used for metabarcoding is expected to exhibit substantial length variation among taxa, the ITS Pipeline Workflow (v1.8) for DADA2 (https://benjjneb.github.io/dada2/ITS_workflow.html) was followed. No length thresholds were applied, with the parameters set as --p-trim-left-f 0, --p-trim- left-r 0, --p-trunc-len-f 0, and --p-trunc-len-r 0. Finally, sequences with fewer than four reads were excluded by applying the filter-feature function with the parameter --p-min-frequency 4.

For the obtained sequences, BLAST 2.13.0+ (Camacho et al. 2009) searches were performed with an E-value threshold of 1e-40. The reference databases used included (1) a new DNA datas of Trichoptera (caddisfly) species from Kamikōchi sequenced for this study (DNA Data Bank of Japan accession numbers: LC859418–LC859545), (2) data retrieved from GenBank (https://blast.ncbi.nlm.nih.gov/Blast.cgi) using the search query “Animalia 16S NOT ’whole genome’ NOT ’chromosome’ NOT ’complete genome’,” which yielded 432,478 sequences (as of January 30, 2025), and (3) the insect DNA database developed by Kanagawa Prefecture (https://www.pref.kanagawa.jp/docs/b4f/suigen/edna-en.html). The following similarity thresholds were applied for taxonomic identification based on BLAST results: ≥98% for the species level, 98–95% for the genus level, 95–90% for the family level, 90–85% for the order level, and 85–80% for the class level.

### 2.4 Running Phylo-MIP

The Phylo-MIP was run using the BLAST search result file as input and the ‘--class’ option to extract ASVs identified as Insecta (80.00 <= pident). Phylogenetic analysis was performed using the neighbor-joining method, and the ‘--bootstrap’ option was applied, with 1000 bootstrap runs performed to estimate node support. This time, the phylogenetic analysis was performed without applying the ‘--treè option, and only data with assigned taxa was output. The output file was checked and some inaccurate data was corrected appropriately. After the taxonomic information was corrected, Phylo-MIP was run on the file with the ‘--onlyp’ option, which performs only phylogenetic analysis and species delimitation analysis. The bPTP analysis was performed using MCMC for 2,000,000 generations, sampling every 200 generations and excluding the first 10% as burn-in.

### 2.5 Similarity between water sampling sites based on Jaccard distance

Based on the presence/absence data of MOTUs across water sampling sites, Jaccard distances were calculated to evaluate community similarity among sites. The calculation of Jaccard distance was performed using the R package vegan version 2.6-8 (Oksanen et al. 2024). Hierarchical clustering based on Ward’s method and the construction of a heatmap were then conducted using the computed Jaccard distance matrix. Hierarchical clustering was performed with the R package dendextend version 1.19.0 (Galili 2015), and the heatmap was generated using the R package pheatmap version 1.0.12 (Kolde 2019).

## 3 Results

### 3.1 Comparison of species lists detected from BLAST and Phylo-MIP analyses

Using eDNA data obtained from six sites in Kanagawa Prefecture in the previous study, we compared species identification results based on general BLAST analysis, which relies on sequence similarity, with those obtained using Phylo-MIP, a phylogenetically based method proposed in this study. Presence/absence data from both approaches were used to assess community similarity through cluster analysis and heatmap visualization (Table 1; Fig. 2, 3, S1). For Phylo-MIP, results from three different methodological variants were also compared. All methods consistently showed high similarity among the three replicates at each site, with the exception of Site 4. Furthermore, no significant differences in community similarity were observed among the methods when comparing across sites. However, the number of detected species was substantially higher with Phylo-MIP. While BLAST identified 127 species, Phylo- MIP detected 162 species (Table 1). This increase was consistent across all six sites (Table 1; Fig. 2).

**Figure 2.**
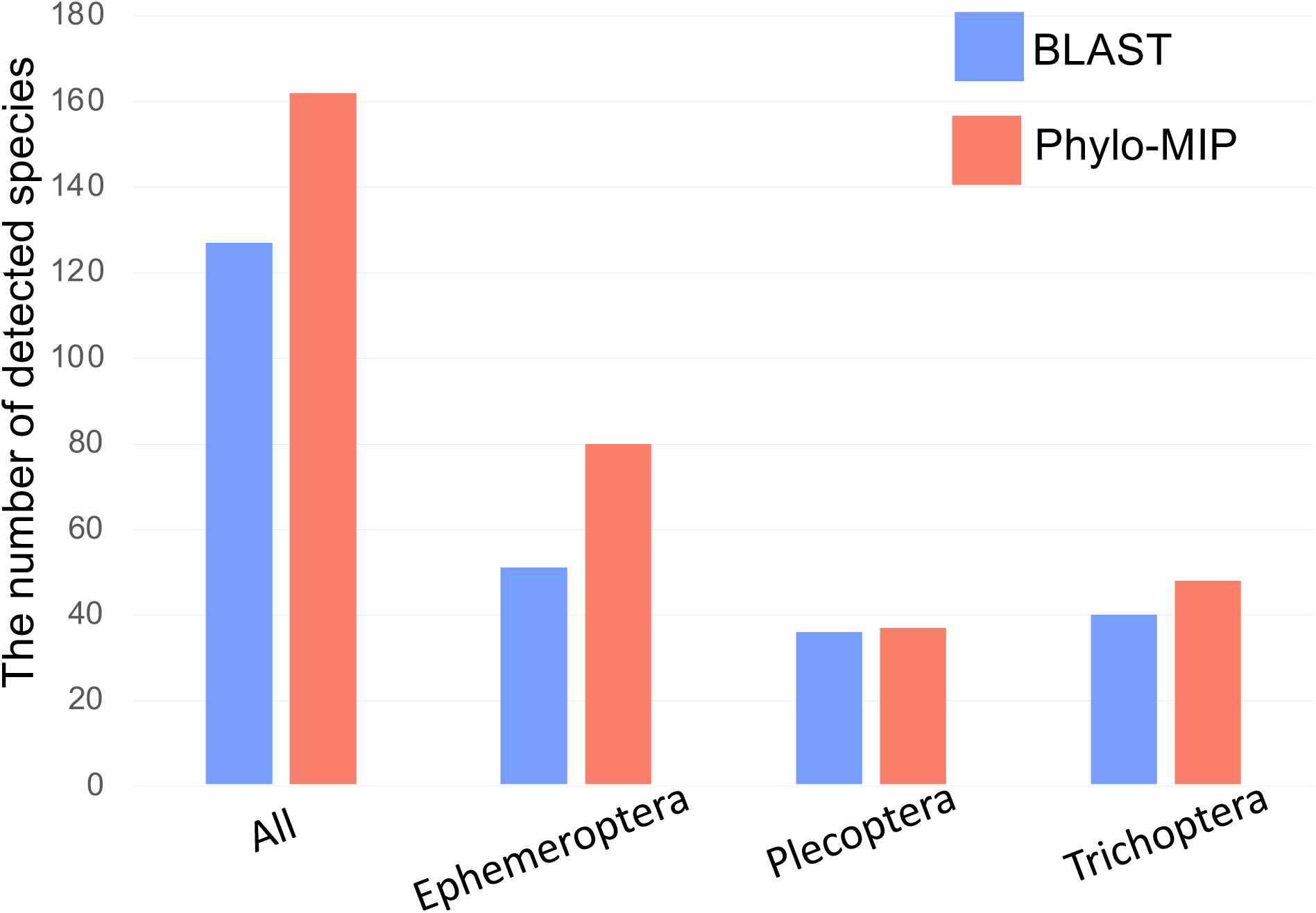
The differences in molecular identification results between BLAST-only search and Phylo-MIP. The y-axis represents the number of detected species, and the x-axis shows taxonomic groups and All.

**Figure 3.**
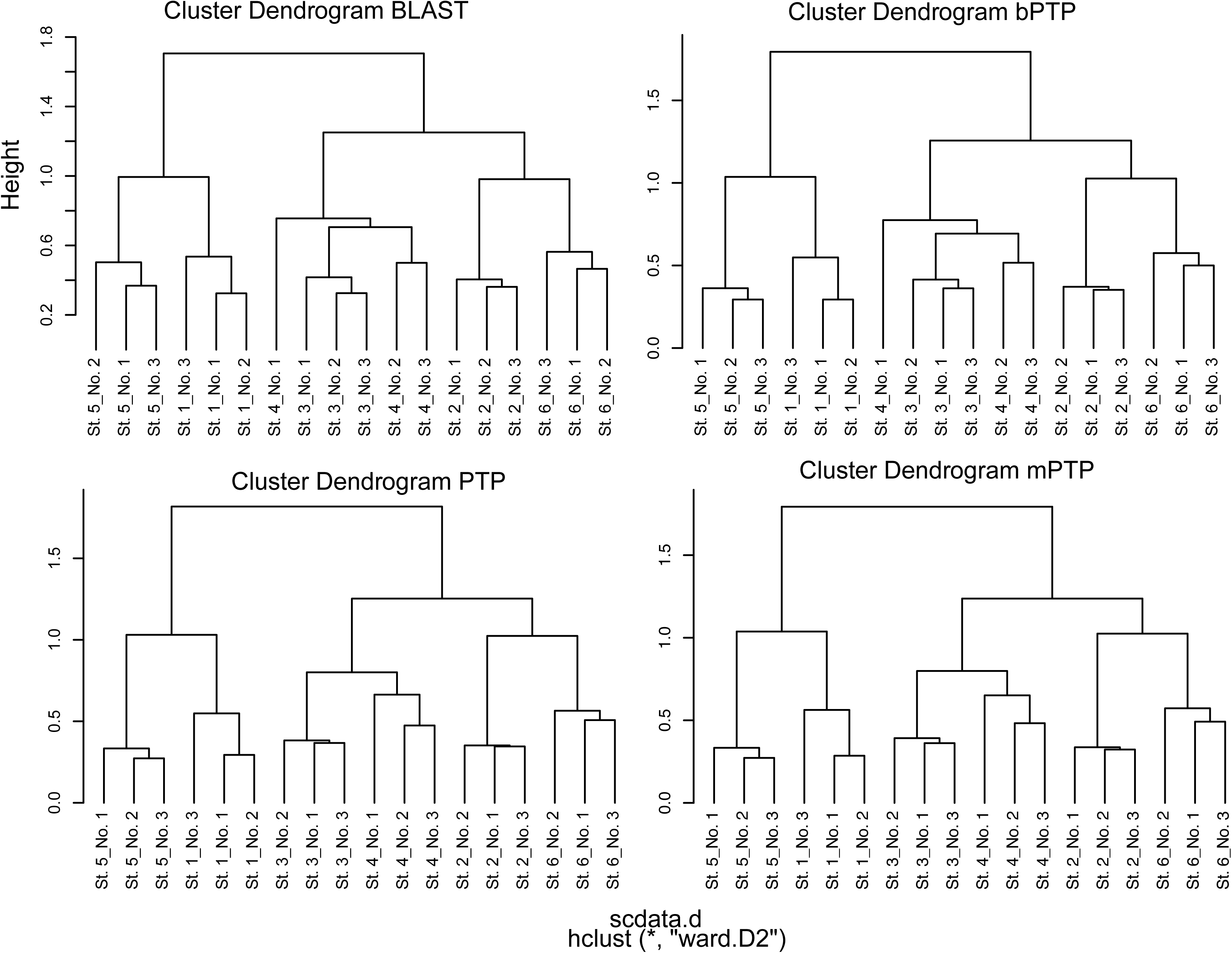
Cluster analysis based on species lists detected by different eDNA metabarcoding approaches in rivers of Kanagawa: BLAST-only and species delimitation methods (bPTP, PTP, mPTP).

**Table 1.** Comparison of the number of detected species by metabarcoding analysis using conventional BLAST-only and Phyla-MIP methods.

| BLAST | St. 1 | St. 2 | St. 3 | St. 4 | St. 5 | St. 6 | all sites |
| --- | --- | --- | --- | --- | --- | --- | --- |
| all | 42 | 84 | 46 | 45 | 34 | 53 | 127 |
| Ephemeroptera | 23 | 29 | 26 | 24 | 22 | 21 | 51 |
| Plecoptera | 2 | 29 | 6 | 9 | 3 | 17 | 36 |
| Trichoptera | 17 | 26 | 14 | 12 | 9 | 15 | 40 |
| Phylo-MIP |  |  |  |  |  |  |  |
| all | 48 | 105 | 52 | 58 | 38 | 68 | 162 |
| Ephemeroptera | 24 | 48 | 32 | 35 | 24 | 34 | 80 |
| Plecoptera | 4 | 28 | 6 | 9 | 3 | 17 | 37 |
| Trichoptera | 20 | 29 | 13 | 14 | 11 | 16 | 48 |

BLAST, Species identification based on sequence similarity using conventional BLAST; Phylo-MIP, Species identification using the method developed in this study

### 3.2 Phylo-MIP DNA metabarcoding at Kamikochi river ecosystem

We conducted water sampling surveys across various environmental at Kamikochi, Nagano Prefecture, and performed eDNA analysis using the Phylo-MIP approach established in this study (Table S2, S3; Fig. 4). The number of reads per sample ranged from 19,779 to 95,454. Based on the presence/absence data of the detected species, community analyses revealed distinct clustering patterns (Fig. 5). Specifically, sites located in relatively fast-flowing environments such as the main stream and tributaries formed separate clusters from those in slower-flowing habitats, including wetlands and small spring streams.

**Figure 4.**
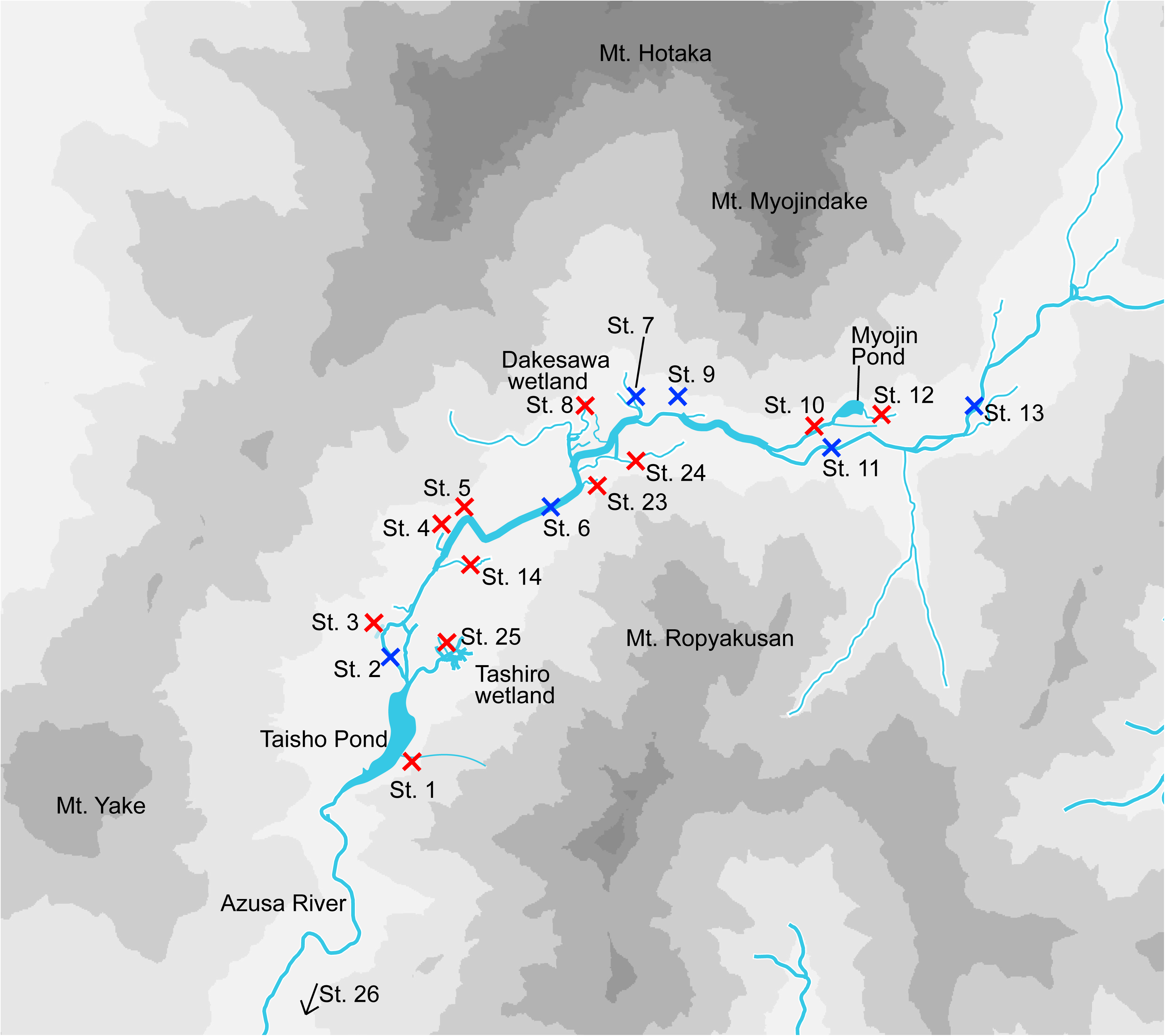
Survey sites in Kamikochi. Black arrows indicate the direction of river flow.

**Figure 5.**
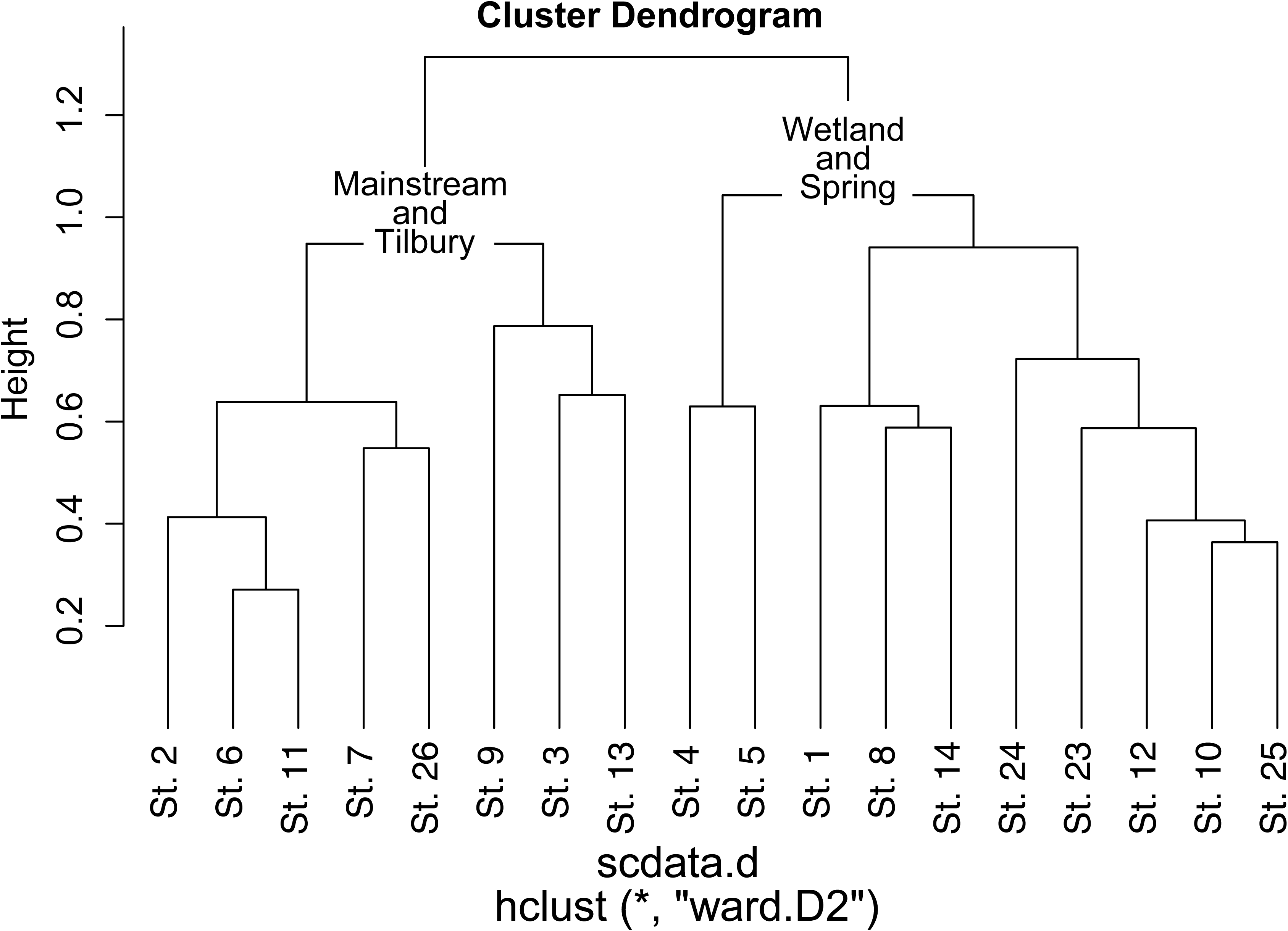
Cluster analysis based on species lists detected by eDNA metabarcoding analysis in Kamikochi.

## 4 Discussion

Environmental DNA metabarcoding is a powerful tool for biodiversity assessment. However, species identification in this context primarily relies on sequence similarity via BLAST. BLAST-based identification is prone to misidentification or failure to identify species due to limitations such as incomplete reference databases, sequence conservation, and low resolution among closely related species (Burian et al. 2021; Keck et al. 2023). This issue is particularly serious in taxonomic groups with high species diversity, such as insects (Takenaka et al. 2025). Several studies have proposed methods that incorporate phylogenetic interpretation to address these issues (Matsen et al., 2010; Barbera et al., 2019; Inoue et al., 2024). In this study, we established a species identification method for DNA metabarcoding that integrates phylogenetic analysis with species delimitation. Phylogenetic information helps mitigate the aforementioned problems and reduces taxon-specific biases caused by sequence similarity thresholds. Furthermore, species delimitation analysis enables objective interpretation of phylogenetic results.

### 4.1 False negatives and Species Delimitation Analysis

Incorporating phylogenetic analysis allows for a more objective interpretation of metabarcoding results, especially in cases where reference databases are incomplete (Matsen et al., 2010; Barbera et al., 2019; Keck et al. 2023; Inoue et al., 2024; Takenaka et al. 2025). Since phylogenetic analysis requires specialized knowledge, we developed an automated pipeline, Phylo-MIP, and made it available on GitHub. False negatives represent a significant obstacle in accurate biodiversity assessment (Burian et al. 2021). These can arise from various factors, including limitations in data analysis. In particular, underrepresentation in reference databases can lead to underestimation of species diversity in metabarcoding analyses (Keck et al. 2023). This not only distorts our understanding of ecosystems and conservation planning but also risks overlooking rare species. By using Phylo-MIP, the number of detected species increased by approximately 28% compared to conventional methods.

This increase is not only due to the detection of sequences that were previously undetectable because of the absence of matching references in databases, but also because sequences that were previously identified only to the genus level could now be resolved to the species level. For example, the genus *Epeorus* (Ephemeroptera) includes multiple species, but when reference data are scarce, sequences are often assigned as *Epeorus* sp. In such cases, even if multiple species are present, they are treated as a single species. However, with Phylo-MIP, phylogenetic analysis allows us to distinguish among genetically distinct lineages such as *Epeorus* sp. 1, sp. 2, and sp. 3, thereby reducing the underestimation of species diversity.

However, this Phylo-MIP approach also presents a concern that must be considered. The PTP-based species delimitation analysis, while designed to delineate species boundaries, can sometimes lead to overestimation of species diversity. Since species are defined based on the degree of genetic divergence, the method identifies MOTUs (Molecular Operational Taxonomic Units), which represent genetically distinct units (Ranasinghe et al. 2022). Traditional biodiversity assessments, however, are typically based on taxonomic species, defined by morphological characteristics. Therefore, discrepancies between genetic species and morphological species must be carefully considered (Frankham et al. 2012; Goldstein et al. 2000). To address this topic, it is necessary to either establish evaluation criteria tailored to DNA metabarcoding or to develop a comprehensive DNA reference database.

In this study, we employed multiple species delimitation methods to avoid overestimation by any single approach. Although not incorporated in the current analysis, future studies should consider integrating barcode gap-based methods such as ABGD (Puillandre et al. 2012) and ASAP (Puillandre et al. 2020). Additionally, to align the results more closely with traditional taxonomic assessments, we treated sequences as belonging to the same species if they were identified as such by either BLAST or species delimitation analysis.

This approach enables more accurate and objective species identification than traditional BLAST-based methods, thereby improving the reliability of biodiversity assessments (Keck et al. 2023). It is especially effective for taxa with insufficient reference data or high species similarity, where the combination of phylogenetic analysis and species delimitation significantly reduces the risk of misidentification or non-identification. Moreover, the automation provided by Phylo-MIP reduces the burden on researchers and enhances reproducibility and efficiency in analysis. In conservation biology and ecosystem monitoring, improved detection accuracy of rare and endangered species facilitates the development of more appropriate conservation strategies. This method has the potential to become a new standard in eDNA metabarcoding and is expected to be widely applied in future biodiversity research.

#### Community Structure of the Freshwater Ecosystem in Kamikochi

In this study, we applied the newly developed pipeline Phylo-MIP, which integrates phylogenetic analysis and species delimitation, to a region where eDNA analysis had not previously been conducted. The target area was Kamikochi, located in Matsumoto City, Nagano Prefecture, Japan. Kamikochi is a unique subalpine zone situated at an altitude of 1500–1600 meters, lying alongside the Azusa River surrounded by mountain ranges exceeding 2500 meters.

Despite its elevation, Kamikochi features a wide and topographically flat area with diverse aquatic habitats, including main stream, meandering streams, tributaries, spring streams, and wetlands (Cited; Milner et al. 2021). Using eDNA metabarcoding analyses, we found that aquatic insect communities differed significantly between large flowing water environments (main and tributary streams) and smaller or stagnant water bodies (springs and wetlands). Aquatic insects are generally specialized to either flowing) or still environments, and species that inhabit both are rare (Larson, 1997; Ribera & Vogler, 2000; Takenaka et al. 2021). This result demonstrates that our method successfully captures habitat-specific biodiversity patterns. In this study, we also used the AQdb-16S primers set that amplifies longer fragments than MtInsects-16S primers set at four survey sites. For data analysis, we used only the homologous regions shared across all samples. Although different primers were used, the community composition was separated by environment rather than by primer type, suggesting that the primers may have similar amplification efficiency. However, since we did not analyze the same samples with both primer sets, further investigation is needed.

We basically used the MtInsects-16S primer, designed for broad coverage of insect taxa (Takenaka et al. 2023). However, the eDNA metabarcoding also detected a wide range of non-insect taxa, including Crustaceans, Cnidarians, etc., (Takenaka et al. 2024), Japanese macaques and fish species (Takenaka et al. 2024; Nagahara et al. 2025). Although we did not compare detection accuracy with taxon-specific primers, the presence of these taxa suggests the primer’s broad utility.

Among freshwater invertebrates, we detected aquatic organisms known to inhabit, e.g., *Physa acuta*, *Euglesa* spp., Annelida, and Platyhelminthes. For fish, Kamikochi is known to inhabiting five species: *Salmo trutta*, *Salvelinus fontinalis*, *Salvelinus japonicus*, and *Rhynchocypris lagowskii*, and *Misgurnus anguillicaudatus* (Peterson et al. 2024). All were successfully detected, along with *Salvelinus leucomaenis*, which may reflect introgressive hybridization among salmonids or limitations in taxonomic resolution (Table S4; Peterson et al. 2024). Additionally, we detected amphibians also inhabited in Kamikochi such as *Onychodactylus* sp. (the sequence showed 97% similarity to *Onychodactylus japonicus*), birds including *Gallinago* sp. (the sequence showed 97% similarity to *Gallinago gallinago*) and *Anas platyrhynchos* (Table S4; Fujiyama et al., 1988; Japan Alps Kamikochi Official website: https://www.kamikochi.or.jp/learn/ecology/birds/, accused at 26 October 2025). Also, terrestrial hexapods such as springtails (Collembola), which are known to inhabit riparian zones (Table S4; Sterzyńska et al., 2014).

## Conclusion

By integrating phylogenetic analysis and species delimitation into the metabarcoding workflow, our method increased the number of detected insect species by approximately 28%. Moreover, it enabled the detection of not only aquatic insects but also amphibians, fish, and riparian terrestrial arthropods inhabiting Kamikochi. Previous studies of eDNA study (Takenaka et al. 2024) have also reported the detection of terrestrial insects, highlighting the high versatility of insect-targeted primers. Notably, aquatic insects tend to dominate eDNA reads due to their high biomass in river systems, with over 90% of reads originating from insects (Takenaka et al. 2024).

In protected areas such as national parks, where conventional surveys may risk disturbing the environment, eDNA analysis offers a non-invasive and effective alternative. By incorporating Phylo-MIP, it becomes possible to assess species diversity more accurately and objectively, minimizing the risk of underestimation. This approach contributes to the generation of reliable and regularly updated monitoring data, which is essential for tracking the distribution of vulnerable, endangered, and invasive species. eDNA-based methods offer substantial advantages, including higher cost- and time-efficiency compared to many traditional survey techniques (Evans et al., 2017), noninvasive monitoring (Cristescu and Hebert, 2018), and high specificity and sensitivity (Wilcox et al., 2013; Leese et al. 2021; Takenaka et al. 2025). The addition of Phylo-MIP further enhances the precision of biodiversity assessments.

This study also demonstrates that sequence data obtained through eDNA metabarcoding can be used not only for similarity-based identification (using only the BLAST) but also for phylogenetic analysis. Phylogenetic interpretation is a critical component in taxonomy, and our approach provides a valuable framework for taxonomic studies. Indeed, we detected numerous potentially undescribed or cryptic species, suggesting that eDNA metabarcoding can serve as a powerful starting point for uncovering hidden biodiversity. As such, this method holds significant potential for advancing taxonomy and biodiversity research.

Ideally, DNA metabarcoding analyses should be conducted with access to a complete and curated reference database (Keck et al. 2023). However, in reality, the expansion of such databases remains a major challenge, particularly for insects, which often exhibit regional genetic differentiation within species (Takenaka & Tojo 2019; Takenaka et al. 2025). Given the large number of undescribed species still being reported, constructing a truly comprehensive database is extremely difficult. In this context, the phylogeny-informed approach established in this study provides an important interpretive framework that helps avoid underestimation of species richness—a common limitation of database-dependent identification methods.

## Acknowledgements

We thank to Prof. Tojo K, Prof. Matsumoto, Ms. Tsuchihashi A, Ms. Nagahara E, Mr. Ichimura K (Shinshu University) for supporting our research. We thank to Ministry of the Environment, agency for cultural affairs, the Forestry Agency, Nagano Prefecture, Matsumoto city for supporting our research. This study was supported by the River Fund of the River Foundation (2021-5311-005, 2022-5311-016 to MT), by a research grant from the Institute of Mountain Science, Shinshu University (2023-903, 2024-1112 to MT), by Nature Positive Sustainable Development Hub of COI-NEXT (JST Grant Number JPMJPF2206).

## Data Accessibility and Benefit-Sharing Genetic data

Raw sequence reads are deposited in the SRA (BioProject: PRJDB38044; BioSample: BioSample: SAMD01706433-SAMD01706450). Unique haplotype data are deposited to NCBI Nucleotide Database (LC859418–LC859545). All methods information is included in this manuscript. All specimens used in this study are stored in the Tojo laboratory of Shinshu University.

## Benefit-Sharing

Benefits Generated: Benefits from this research accrue from the sharing of our data and results on public databases as described above and a research collaboration was developed with scientists from the countries providing genetic samples, all collaborators are included as co-authors

## Author Contributions

Masaki Takenaka: designed research (lead); performed research (lead); contributed analytical tools (supporting); analyzed data (supporting); wrote the paper (lead); review and editing (lead). Takumi Yoshida: performed research (lead); contributed analytical tools (lead); analyzed data (lead); wrote the paper (lead); review and editing (supporting). Shonosuke Shigeta: contributed analytical tools (lead); analyzed data (lead); wrote the paper (lead); review and editing (supporting). Yuta Hasebe: analyzed data (lead); review and editing (supporting).

## Supporting/Supplemental Information

Table S1 The number of reads detected by eDNA analysis for Insecta in the Kanagawa River. Table S2 Water sampling sites in Kamikochi.

Table S3 Species list detected by eDNA analysis within Insecta at Kamikochi.

Table S4. The number of reads detected by eDNA analysis for non-insect invertebrates at Kamikochi

---

**Figure S1.**
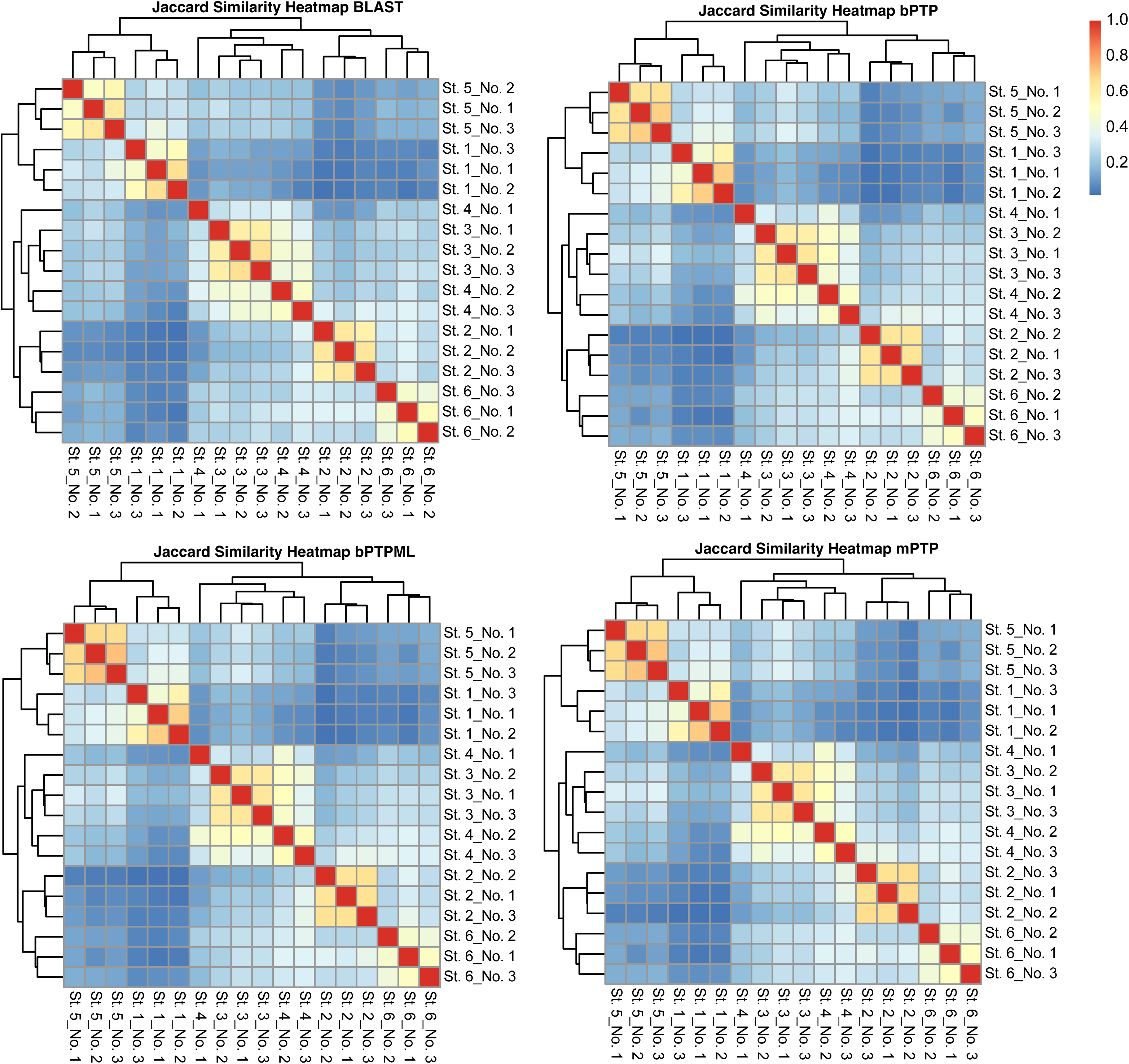
Jaccard Similarity Heatmap based on species lists detected by different eDNA metabarcoding approaches in rivers of Kanagawa: BLAST-only and species delimitation methods (bPTP, PTP, mPTP).

